# The IgG3 Subclass of β1-adrenergic receptor autoantibody is an endogenous biaser of β1AR signaling

**DOI:** 10.1101/2020.09.17.302059

**Authors:** Maradumane L. Mohan, Yuji Nagatomo, Sromona D. Mukherjee, Timothy Engelman, Rommel Morales, W.H. Wilson Tang, Sathyamangla V. Naga Prasad

**Affiliations:** Department of Cardiovascular and Metabolic Sciences, Lerner Research Institute, Cleveland Clinic, Cleveland, OH, 44195; Department of Cardiology, National Defense Medical College, Tokorozawa, Japan; Heart and Vascular Institute, Cleveland Clinic, Cleveland, OH

## Abstract

Autoantibodies recognizing human β1ARs generated due to dysregulation in autoimmune response are generally associated with deleterious cardiac outcomes. However, cellular studies show that isolates of β1AR autoantibody from patients differentially modulate β1AR function. β1AR autoantibodies belong to the IgG class of immunoglobulins, however it is not known whether the IgG sub-classes mediate variability in β1AR responses. To determine whether the IgG3 subclass of β1AR autoantibodies uniquely modulate β1AR function, HEK293 cells stably expressing human β1ARs were utilized. Treatment of cells with IgG3(-) serum resulted in significant increase of cAMP compared to IgG3(+) serum. Pre-treatment of cells with IgG3(+) serum impaired dobutamine-mediated Adenylate Cyclase (AC) activity and cAMP generation whereas, it surprisingly increased AC activity and cAMP generation with β-blocker metoprolol. Consistently, purified IgG3(+) β1AR autoantibodies impaired dobutamine-mediated cAMP while elevating metoprolol-mediated AC activity and cAMP. Despite IgG3(+) autoantibodies reducing cAMP response to dobutamine, they mediate significant ERK activation upon dobutamine. IgG3(+) β1AR autoantibodies did not alter β2AR function, reflecting their specificity. The study shows that IgG3(+) β1AR autoantibody impairs agonist-mediated G-protein coupling while preferentially mediating G-protein-independent ERK activation. Furthermore, it uniquely biases β-blocker towards G-protein coupling. This unique biasing capabilities of IgG3(+) β1AR autoantibodies may underlie the beneficial outcomes in patients.

## INTRODUCTION

β-adrenergic receptors (βARs) are one of the most well studied proto-typical 7-trans-membrane receptors that are powerful regulators of cardiac function (Rockman et al., 2002; Vasudevan et al., 2011a). Among the βARs, β1- and β2-ARs are highly expressed in the myocardium (Rockman et al., 2002). Catecholamine binding to the βARs results in G-protein coupling leading to adenylyl cyclase activation, mediating cAMP-protein kinase A (PKA) signal cascade (Wallukat, 2002). Activation of cAMP-PKA cascade alters calcium cycling resulting in increased myocardial contractility. Although β1- and β2-ARs play a key role in myocardial contraction, there is also increasing appreciation that beyond contraction, they may have distinct roles to play in phenotypic outcomes (Dungen et al., 2020; Liaudet et al., 2014). Increasing evidence from in vivo and in vitro studies show functional divergence between β1AR and β2ARs. Cellular studies show that chronic catecholamine stimulation leads to cardiomyocyte apoptosis through activation of β1ARs, while β2ARs may mediate cardioprotective signaling (Milano et al., 1994; Steinberg, 2018). Consistently, cardiomyocyte overexpression of β1ARs leads to maladaptive cardiac remodeling, while overexpression of β2ARs results in hypertrophic response, reflecting their divergent roles (Engelhardt et al., 1999; Steinberg, 2018). However, β1AR downregulation (loss of cell surface receptors) and desensitization (inability to be activated by catecholamine) are one of the key hallmark features of human heart failure (Port and Bristow, 2001; Steinberg, 2018).

Dilated cardiomyopathy (DCM) is one the most commonly observed phenotype of heart failure associated with progressive loss in ventricular function, wherein idiopathic DCM represents pathogenesis without a specific known cause (Magnusson et al., 1994; Wynne, 1988). Patients with idiopathic DCM have a diagnosis of diverse etiologies and varied presentations however, dysregulation of the immune system is considered to be one of the central players in cardiac pathogenesis. The human immune system is a complex multicellular regulated defense mechanism that is characterized by inter-individual variability in response to a similar stress/injury. In healthy homeostatic conditions, it is designed to discriminate between self and foreign components, and clear components deemed to be foreign (Crampton et al., 2010; Kaya et al., 2012; Mann, 2011). However, when this regulatory control is lost, it leads to pathological circumstances wherein self-components are attacked, resulting in auto-immune disease (Crampton et al., 2010; Kaya et al., 2012; Mann, 2011). Thus, autoantibodies generated against the self-antigens exacerbate and may accelerate the disease progression (Crampton et al., 2010; Kaya et al., 2012; Mann, 2011).

Circulating autoantibodies have been identified in heart failure and increasing evidence suggests that they may be critically linked to heart failure pathogenesis. Autoantibodies against β1AR have been observed in patients with heart failure (30-40%), and are positively co-related to heart failure (Baba, 2010; Iwata et al., 2001; Jahns et al., 1999b; Magnusson et al., 1990; Magnusson et al., 1994; Nagatomo et al., 2009; Stork et al., 2006). Studies have shown that β1AR autoantibody stabilizes the receptor in an active form prolonging its activation mimicking catecholamines (Deubner et al., 2010). Furthermore, β1AR autoantibodies can elevate the L-type Ca^2+^ current increasing in vitro contractility (Christ et al., 2001). This elevated and prolonged activation reflects the hyper-sympathetic state associated with deleterious cardiac remodeling and DCM. Although β-blockers ameliorate the signaling from the sympathetic overdrive, their role in upregulation of βARs is thought to underlie the worsening outcomes of heart failure due to binding by β1AR autoantibodies. However, recent studies have demonstrated myocardial recovery in patients who are on β-blockers and harbor β1AR autoantibodies belonging to the IgG3 subclass (Nagatomo et al., 2017). β1AR autoantibodies belongs to IgG class of immunoglobulins that can be further sub-classified into IgG 1, 2, 3 and 4 (Schur, 1988; Vidarsson et al., 2014). Therefore, our studies in the current manuscript determine the role of the IgG3 subclass of β1AR autoantibodies in modulating β1AR function/signaling. Serum from patients containing IgG3 -positive [IgG3(+)] and -negative [IgG3(-)], and purified β1AR autoantibodies were used in cellular studies. HEK 293 cells stably expressing human β1AR were pre-treated with β1AR autoantibodies followed by β1AR-selective agonist (dobutamine) or antagonist/blocker (metoprolol). Our studies show that the IgG3 subclass of β1AR autoantibodies markedly attenuates agonist mediated G-protein signaling, while surprisingly biasing the β-blocker signal towards G-protein coupling, unraveling a unique signaling role for this sub-class of β1AR autoantibodies.

## RESULTS

### Human Embryonic Kidney 293 (HEK 293) cells stably expressing human β1AR

To understand how the IgG3 subclass of β1AR autoantibodies modulate the function of human β1AR, we generated the HEK 293 cell line stably expressing FLAG-tagged human β1AR (β1AR-HEK 293 cells). The expression of the FLAG-β1AR in the cells was tested by western immunoblotting using anti-FLAG antibody. Immunoblotting of cell lysates showed robust expression of FLAG-β1AR in β1AR-HEK 293 cells [**Fig. 1A**]. Cell surface expression of the β1ARs was confirmed by radio-ligand binding using _[125]_I-Cyanopindolol on plasma membranes isolated from β1AR-HEK 293 cells. The binding curve showed high levels of β1AR expression on the cell surface [**Fig. 1B**]. To determine whether overexpressed β1ARs were functional, cAMP generation was measured in β1AR-HEK 293 cells treated with either the β1AR-specific agonist dobutamine (Dob) or -blocker metoprolol (Meto). The cAMP generation was significantly increased in cells treated with Dob and was dramatically suppressed with Meto, reflecting functional fidelity of overexpressed FLAG-β1ARs [**Fig. 1C**]. Given that G-protein coupling is a key tenet of β1AR function, we measured G-protein coupling by performing adenylyl cyclase (AC) assay using plasma membrane fractions. Dob treatment significantly increased AC activity which was markedly suppressed with Meto [**Fig. 1D**]. Sodium fluoride (NaF) was used as positive control to directly stimulate AC and to show the integrity of the G-protein coupling complex in the isolated plasma membranes [**Fig. 1D**]. To further test for activation of downstream signaling, β1AR-HEK 293 cell lysates were immunoblotted for phospho-ERK (pERK) levels [**Fig. 1E**]. ERK phosphorylation was significantly increased by Dob stimulation and Meto treatment did not appreciably alter ERK phosphorylation [**Fig. 1F**]. This human β1AR-overexpressing, HEK 293 cell line now provided the primer to determine the role of the IgG3 subclass of β1AR autoantibodies in modulating β1AR function.

**Fig 1:**
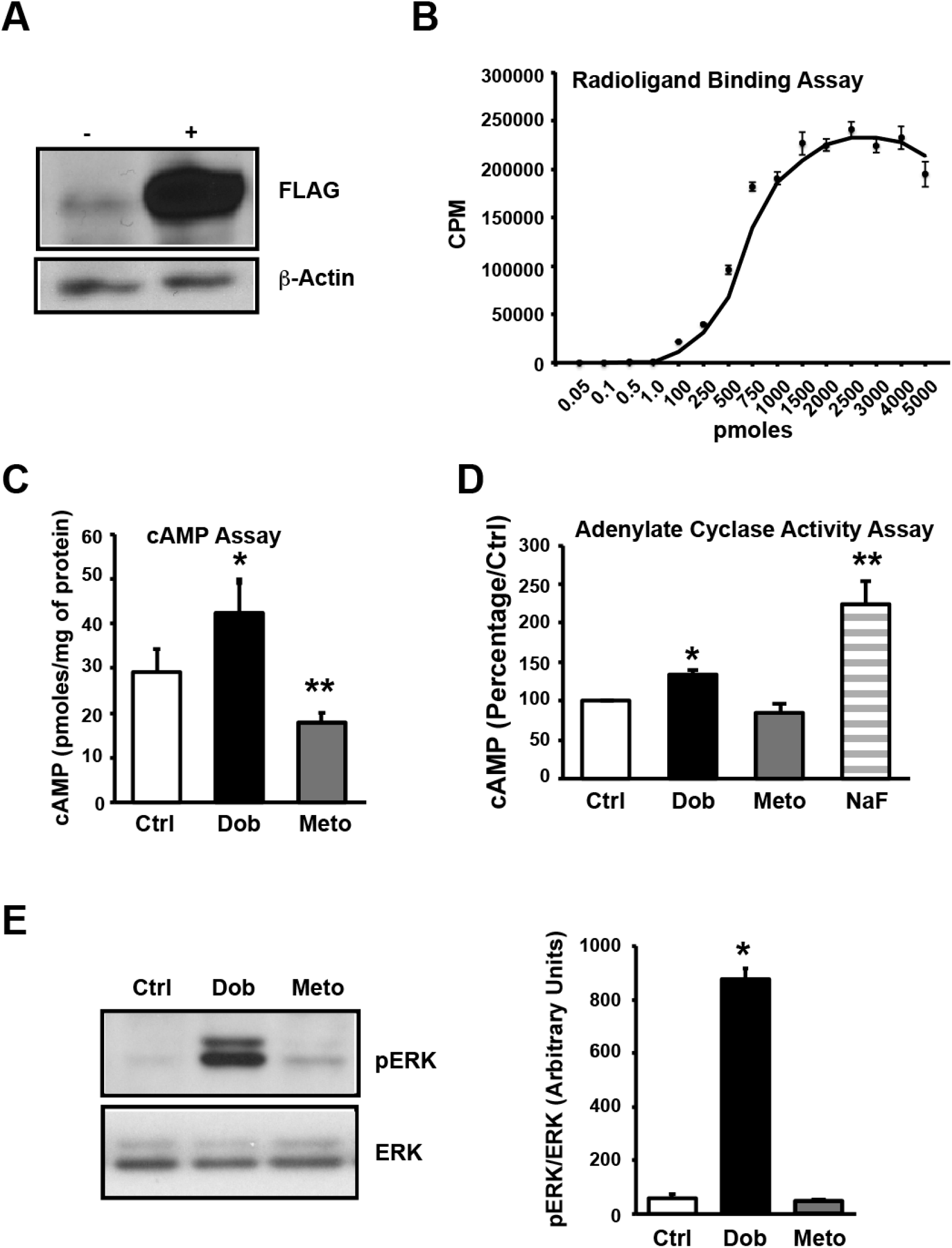
Generation and characterization of stable Human Embryonic Kidney (HEK) cell line expressing human β1-Adrenergic Receptors (β1AR). **(A)** Parental HEK293 cells and HEK293 cells overexpressing FLAG-human-β1AR (HEK-β1AR) were lysed with NP-40 lysis buffer, cell lysates (50 μg each) were subjected to SDS-PAGE and immunoblotted with anti-FLAG antibody. The blots were stripped and immunoblotted with anti-β-actin antibody as loading control. **(B)** Isolated plasma membranes from HEK-β1AR cells were used to perform the receptor binding assay using _125_I -Cyanopindolol to show the expression of receptors on the cell membranes and binding curve generated establish very high expression of the receptors (n=3). **(C)** HEK-β1AR cells were serum starved for 4 h and stimulated with 10 μM Dobutamine (Dob) or 10 μM Metoprolol (Meto) for 10 min. The cells were lysed and cyclic Adenosine Mono Phosphate (cAMP) generation was measured using cAMP assay kit. Bar graphs represent cumulative data (n=3). *p ≤ 0.05 Ctrl vs Dob, **p ≤ 0.01 Meto vs Dob/Ctrl. **(D**) Isolated membranes from HEK-β1AR cells were used to perform Adenylate Cyclase (AC) assay in vitro to assess the function of the expressed receptors in the presence of Dob or Meto. The amount of cAMP generated by control is expressed as 100 percent and percent change in generation of cAMP in treatments are shown (n=3). *p ≤ 0.05 Ctrl vs Dob, **p ≤ 0.001 NaF vs Ctrl/Dob/Meto. **(E)** HEK-β1AR cells were serum starved and stimulated with Dob or Meto. The cell lysates were subjected to western immunoblotting with anti-phospho-ERK. The blots were stripped and immunoblotted with anti-ERK antibody as loading control (left panel). Cumulative densitometric data are presented as bar graphs (n=3). *p ≤ 0.0001, Dob vs Ctrl/Meto (right panel).

### Role of β1AR autoantibodies on cAMP generation

To assess whether β1AR autoantibodies modulate β1AR function, we used the validated β1AR-HEK 293 cells for these studies. Human sera positive for IgG were collected and classified into IgG3(-) and IgG3(+) (classification described elsewhere (Iwata et al., 2001; Nagatomo et al., 2011; Nagatomo et al., 2016; Nagatomo et al., 2017; Nagatomo et al., 2009)). The β1AR-HEK 293 cells were treated with these sera and the cAMP generation measured. Treatment of the cells with IgG3(-) sera significantly increased cAMP whereas, IgG3(+) sera did not alter cAMP levels compared to untreated controls [**Fig. 2A**]. To test whether IgG3-specific sera containing β1AR autoantibody modulates β1AR responses to Dob or Meto treatment, cells were pre-treated with IgG3(-) or IgG3(+) sera, and cAMP generation was measured following Dob or Meto treatment. Pre-treatment of cells with IgG3(-) sera blunted the Dob-mediated cAMP generation, while Meto significantly decreased cAMP production [**Fig. 2B**]. Similarly, pre-treatment with IgG(+) sera blocked cAMP generation following Dob [**Fig. 2C**]. However, surprisingly Meto treatment resulted in significant increase in cAMP generation in IgG3(+) sera treated cells [**Fig. 2C**].

**Fig 2:**
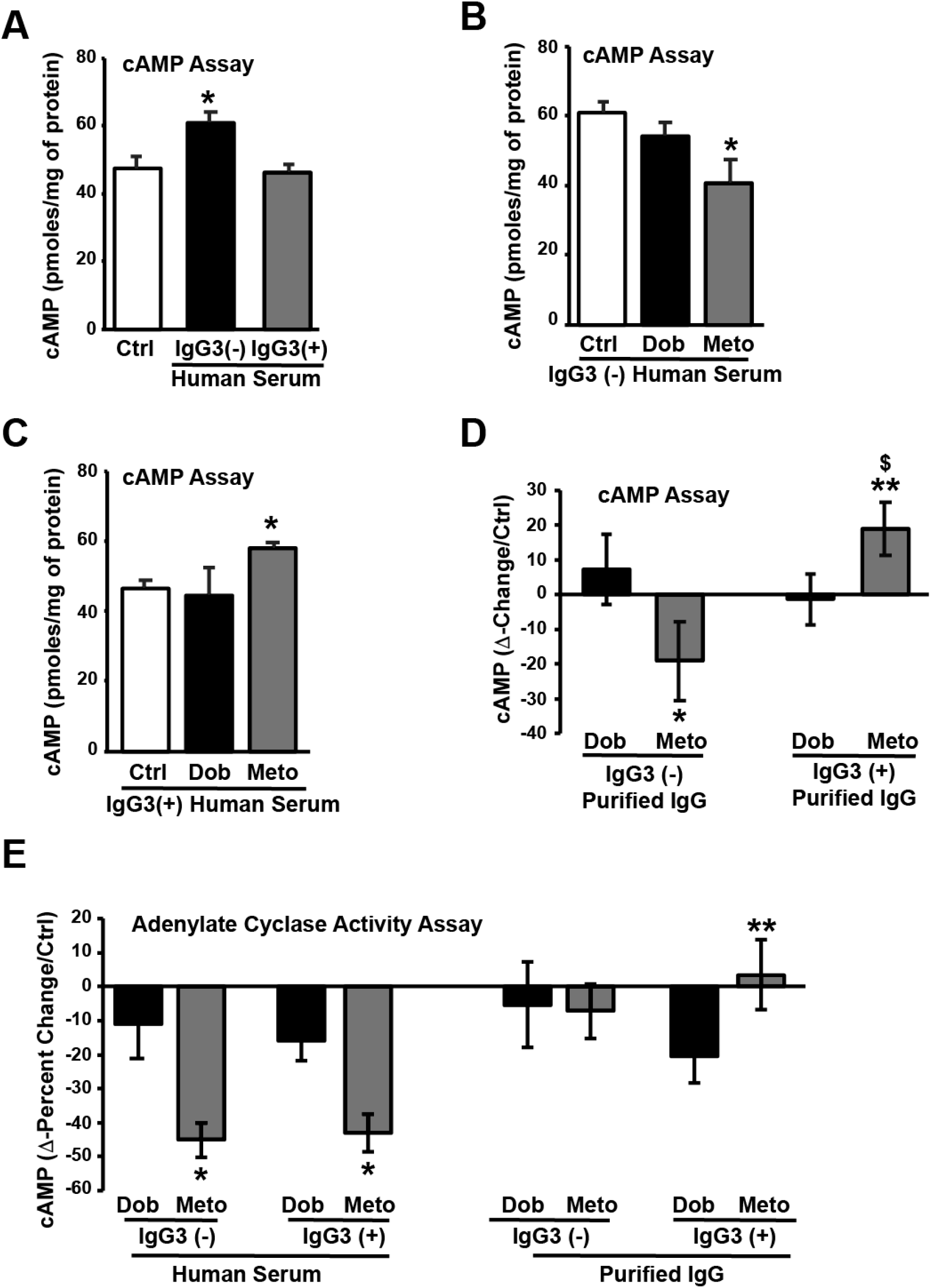
Autoantibodies (AAb) alter β1AR function. **(A)** HEK-β1AR cells were serum starved treated with human serum negative for IgG3 class of AAb [IgG3(-)] or positive for IgG3 class of AAb [IgG3(+)] for 10 min. The cAMP was measured (n=3,4). *p ≤ 0.01 IgG3(-) vs Ctrl/IgG3(+). **(B)** HEK-β1AR cells were serum starved, pre-treated with IgG3(-) human serum for 30 min and stimulated with Dob or Meto. The cAMP was measured (n=3). *p ≤ 0.05 Meto vs Ctrl. **(C)** HEK-β1AR cells were serum starved, pre-treated with IgG3(+) human serum and stimulated with Dob or Meto. The cAMP was measured (n=4). *p ≤ 0.05 Meto vs Ctrl/Dob. **(D)** HEK-β1AR cells were serum starved, pre-treated with affinity purified IgG3(-) AAb for 30 min and stimulated with Dob or Meto. The cAMP was measured (n=9). Δ-change compared to control is represented. *p ≤ 0.05 Meto vs Dob [IgG3(-)]. **p ≤ 0.05 Meto vs Dob [IgG3(+)]. $p ≤ 0.01 Meto [IgG3(+)] vs Meto [IgG3(-)]. **(E)** Isolated membranes from HEK-β1AR cells were pre-treated with IgG3(-) human serum, IgG3(-) purified IgG, IgG3(+) human serum, or IgG3(+) purified IgG, AC activity was performed in the presence of Dob or Meto. The amount of cAMP generated by control is expressed as 100 percent, percent change in generation of cAMP in treatments were calculated and Δ-change compared to control is represented (n=4, 5, 7). *p ≤ 0.01 Meto human serum vs Dob human serum. **p≤ 0.05 Meto IgG3(+) purified IgG vs Dob IgG3(+) purified IgG.

Since sera could contain factors that may potentially alter the regulation of β1ARs by β1AR autoantibodies, the autoantibodies were affinity-purified and used to determine their role in modulating Dob- or Meto-mediated signaling. Interestingly, treatment of cells with purified immunoglobulins, IgG3(-) or IgG3(+) did not alter cAMP generation [**Fig. S1A**] in contrast to the sera treatment [**Fig. 2A**]. Pre-treatment of cells with affinity purified IgG3(-) immunoglobulins resulted in a slight increase in Dob-mediated cAMP generation and significant decrease in cAMP with Meto [**Fig. 2D**], consistent with IgG(-) sera [**Fig. 2B**]. Interestingly, Dob-mediated cAMP generation was abrogated in cells following pre-treatment with affinity purified IgG3(+) immunoglobulins [**Fig. 2D**] suggesting inhibition of G-protein coupling upon agonist binding. Surprisingly, Meto treatment reversed this inhibition resulting in significant generation of cAMP [**Fig. 2D**]. These observations show that both IgG3(-) and IgG3(+) samples have divergent effects with both Dob-mediated cAMP generation as well as Meto-mediated cAMP generation. Thus, IgG3(+) immunoglobulins uniquely modulate β1AR blocker metoprolol to engage in G-protein pathways while contrastingly blocking the classical coupling of G-protein by β1AR agonist dobutamine.

### Assessment of adenylate cyclase activity; A measure of G-protein coupling

To further elucidate how IgG3(+) β1AR autoantibodies allow for differential G-protein coupling, effect of Dob or Meto on AC activity was assessed using isolated plasma membrane in vitro. Pre-treatment of plasma membrane with human serum containing IgG3(-) decreased Dob-mediated AC activity but did not reach significance and significantly reduced AC activity by Meto treatment [**Fig. 2E; Fig. S1D**]. On the contrary, Dob treatment in the presence of purified IgG3(-) autoantibodies increased AC activity but did not reach significance [**Fig 2E; Fig. S1D**]. Pre-treatment with human serum containing IgG3(-) significantly decreased AC activity with Meto treatment. However, purified IgG3(-) immunoglobulins significantly increased AC activity with Meto treatment [**Fig 2E; Fig. S1D**]. This is in contrast to cAMP data wherein pre-treatment with IgG3(-) β1AR autoantibodies resulted in a modest Dob-mediated change in cAMP that did not reach significance, while cAMP levels were significantly reduced in the presence of Meto [**Fig. 2D**]. In contrast, pre-treatment of plasma membranes with human serum containing IgG3(+) and purified IgG3(+) immunoglobulins blocked the classical Dob-mediated increase in adenylyl cyclase activity [**Fig. 2E; Fig. S2E**]. However, Meto treatment in presence of purified IgG3(+) β1AR autoantibodies showed significant restoration of adenylyl cyclase activity **[Fig. 2E; Fig. S2E]** when compared to the plasma membranes that were pretreated with human serum containing IgG3(+) autoantibodies. The data is in consistent with increase in cAMP generation following treatment with IgG3(+) human serum [**Fig. 2C**] and affinity purified IgG3(+) immunoglobulins [**Fig. 2D**]. Thus, these observations strengthen the findings that IgG3(+) β1AR autoantibodies block agonist Dob mediated G-protein coupling while surprisingly allowing Meto to mediate signals through engagement of G-protein coupled pathways.

### IgG3(+) β1AR autoantibodies modulate downstream ERK activation

Activation of ERK is one of the key measures of biased downstream G-protein independent signaling that is mediated by β-arrestins (Shenoy et al., 2006). Given the unique modulation of β1AR by the IgG3(+) β1AR autoantibodies that blocks G-protein coupling upon Dob, we assessed for changes in phospho-ERK following IgG3(-) or IgG3(+) sera and affinity purified antibodies. IgG3(+) serum pre-treatment by itself resulted in significant elevation in phospho-ERK however, Dob or Meto treatment of cells in the presence of IgG3(-) or IgG3(+) sera showed no significant differences in ERK activation [**Fig. 3 A, B and C**]. These observations suggest that both IgG3(-) or IgG3(+) sera impairs Dob mediated ERK activation while Meto does not alter phospho-ERK status. Since sera may contain components that could alter β1AR function and signaling, we tested for ERK activation using affinity purified β1AR autoantibodies. Surprisingly Dob-mediated ERK phosphorylation was restored in cells pre-treated with affinity purified IgG3(-) or IgG3(+) β1AR autoantibodies [**Fig. 3 D, E & F]**. While, Meto treatment did not activate ERK in presence of IgG3(-) or IgG3(+) affinity purified β1AR autoantibodies [**Fig. 3 D, E & F**]. It is important to note that IgG3(+) autoantibodies that completely abrogate G-protein coupling with Dob [**Fig. 2H**] now allows for efficient activation of ERK. This suggests that IgG3(+) autoantibodies could uniquely bias the agonist signaling towards G-protein-independent pathways showing an yet unappreciated role of IgG3(+) autoantibodies in regulating receptor function.

**Fig 3:**
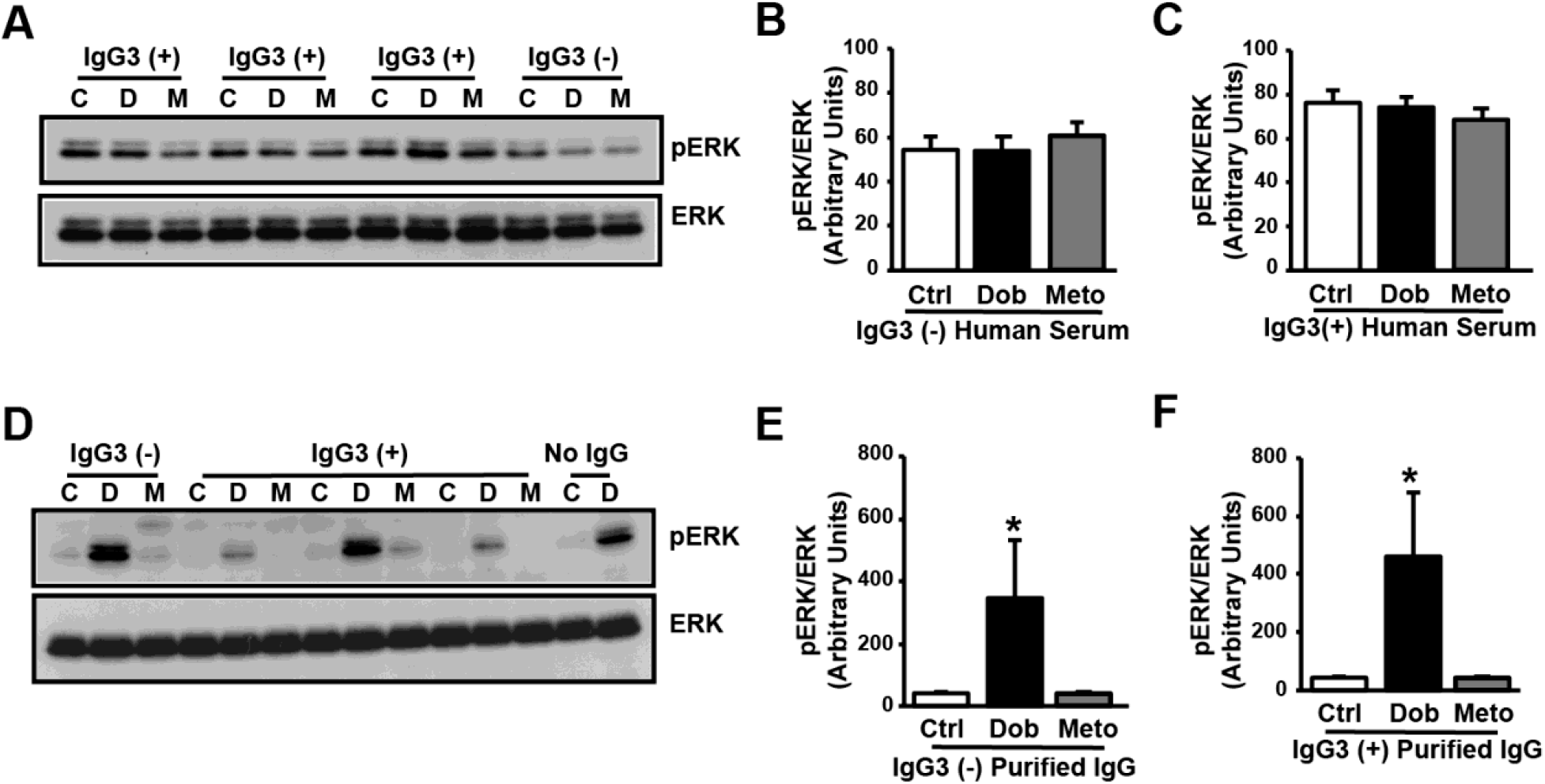
Autoantibodies (AAb) positive human serum alter β1AR signaling. **(A)** HEK-β1AR cells were serum starved for 4 h, pre-treated with IgG3(-) or IgG3(+) human serum for 30 min and stimulated with Dob or Meto. The cell lysates were subjected to western immunoblotting with anti-phospho-ERK antibody. The blots were stripped and immunoblotted with anti-ERK antibody as loading control. **(B)** Cumulative data for cells pre-treated with IgG3(-) human serum (n=3). **(C)** Cumulative data for cells pre-treated with IgG3(+) human serum (n=3). **(D)** HEK-β1AR cells were serum starved, pre-treated with affinity purified IgG3(-) or IgG3(+) AAb and stimulated with Dob or Meto. The cell lysates were subjected to western immunoblotting as above. **(E)** Cumulative data for cells pre-treated with affinity purified IgG3(-) AAb (n=5). *p ≤ 0.0001 Dob vs Ctrl/Meto. **(F)** Cumulative data for cells pre-treated with affinity purified IgG3(+) Aab (n=6, 7). *p ≤ 0.0001 Dob vs Ctrl/Meto.

### β1AR autoantibodies do not alter β2AR signaling

To test whether β1AR autoantibodies are specific to only modulating β1AR signaling, we used HEK 293 cells stably expressing β2AR (β2AR-HEK 293 cells) [**Supplemental Fig S1**) (Shenoy et al., 2006; Vasudevan et al., 2011b; Vasudevan et al., 2013). β2AR-HEK 293 cells were pretreated with either no sera, IgG3(-) or IgG(+) sera. Following pre-treatment the cells were stimulated with isoproterenol (ISO, βAR agonist) or ICI (an inverse β2AR antagonist) and phospho-β2AR levels were assessed by immunoblotting. There was significant increase in β2AR phosphorylation following ISO treatment with either no sera, IgG3(-) or IgG(+) sera [**Fig. 4 A, B, C & D**]. Similarly, IgG(-) or IgG(+) sera did not alter phosphorylation state of β2ARs upon ICI treatment [**Fig. 4 A upper panel, B, C & D**]. These studies show that IgG3(-) or IgG(+) sera do not alter β2AR responses to ISO. FLAG immunoblotting was performed as loading control [**Fig. 4A lower panel**]. To further validate that IgG(-) or IgG(+) sera does not alter downstream signaling, phospho-ERK was assessed following pre-treatment with sera and stimulation with ISO or ICI. Pre-treatment with either IgG3(-) or IgG3(+) sera did not affect ISO-mediated ERK activation or alter ICI dependent responses [**Fig. 4 E, F, G & H**]. Together these data suggest that sera containing β1AR autoantibodies from either the IgG3(-) or IgG3(+) family of immunoglobulins does not affect the activation/downstream signaling of β2ARs reflecting the specificity of these autoantibody regulation.

**Fig 4:**
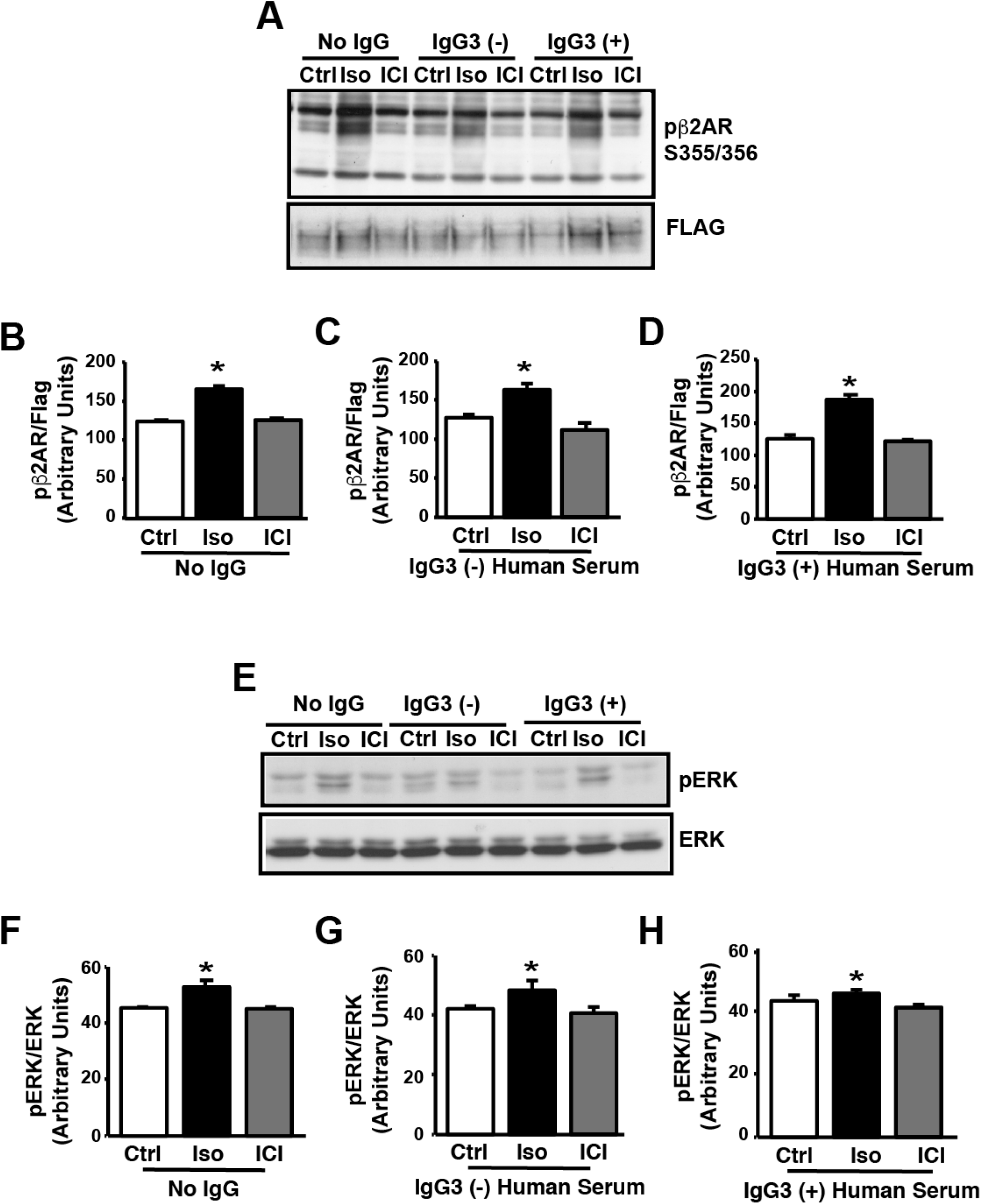
Autoantibodies (AAb) positive human serum does not alter β2AR signaling. **(A)** HEK293 cells overexpressing FLAG-human-β2AR (HEK-β2AR) were serum starved for 4 h, pre-treated with no IgG, IgG3(-) or IgG3(+) human serum for 30 min and stimulated with 10 μM Isoproterenol (Iso) or 10 μM ICI (ICI) for 10 min lysed with NP-40 lysis buffer, cell lysates (50 μg each) were subjected to SDS-PAGE and immunoblotted with anti-phospho-β2AR antibody. The blots were stripped and immunoblotted with anti-FLAG antibody as loading control. The cell lysates were subjected to western immunoblotting with anti-phospho-β2AR. The blots were stripped and immunoblotted with anti-FLAG antibody as loading control. **(B)** Cumulative data for cells pre-treated with no IgG3 human serum (n=3). *p ≤ 0.01 Iso vs Ctrl/ICI **(C)** HEK-β2AR cells were serum starved, pre-treated with IgG3(-) human serum and stimulated with Iso or ICI. The cell lysates were subjected to western immunoblotting as above (n=3). *p ≤ 0.01 Iso vs Ctrl/ICI. **(D)** HEK-β2AR cells were serum starved, pre-treated with IgG3(+) human serum and stimulated with Iso or ICI. The cell lysates were subjected to western immunoblotting as above (n=4). *p ≤ 0.01 Iso vs Ctrl/ICI. **(E)** HEK-β2AR cells were serum starved, pre-treated with no IgG, IgG3(-) or IgG3(+) human serum and stimulated with Iso and ICI. The cell lysates were subjected to western immunoblotting with anti-phospho-ERK. The blots were stripped and immunoblotted with anti-ERK antibody as loading control. **(F)** Cumulative data for cells pre-treated with no IgG human serum (n=3). *p ≤ 0.01 Iso vs Ctrl/ICI. **(G)** Cumulative data for cells pre-treated with IgG3(-) human serum (n=3). *p ≤ 0.05 Iso vs Ctrl/ICI. **(H)** Cumulative data for cells pre-treated with IgG3(+) human serum (n=4). *p ≤ 0.05 Iso vs Ctrl/ICI.

## DISCUSSION

β1AR autoantibodies have been found in 30 – 95% of patients with a diagnosis of idiopathic dilated cardiomyopathy (DCM) (Limas et al., 1992; Wallukat et al., 1991) and higher percentages are consistently associated with patients requiring left ventricular assist device (LVAD) (Dandel et al., 2012; Youker et al., 2014). Such a co-relation suggests a role for β1AR autoantibodies in the progression of heart failure. However, recent studies on patients enrolled in the IMAC-2 (Intervention in Myocarditis and Acute Cardiomyopathy-2) study found favorable myocardial outcomes in patients with β1AR autoantibodies belonging to the IgG3 subclass (Nagatomo et al., 2016; Nagatomo et al., 2017). This suggests the IgG3 subclass of β1AR autoantibodies may uniquely regulate β1AR function, providing benefits compared to the known deleterious role of the IgG class of β1AR autoantibodies. Using IgG3(-) or IgG3(+) β1AR autoantibody-containing sera and affinity-purified β1AR autoantibodies, we show that the IgG3(+) subclass of the antibodies blunt β1AR response to the β1AR agonist, dobutamine. However, the IgG3(+) subclass of β1AR autoantibodies modulates the β1AR-blocker, metaprolol-dependent signaling. Surprisingly IgG3(+) β1AR autoantibodies increases adenylyl cyclase activity and cAMP levels with metoprolol treatment, suggesting a yet, unappreciated role of the IgG3(+) subclass of β1AR autoantibodies that could now bias the metoprolol signal endogenously. This unique signaling mechanism may potentially underlie the beneficial outcomes observed in patients.

Autoantibodies classically belong to the IgG class of immunoglobulins which are one of the abundant among the five classes (IgM, IgE, IgG, IgA and IgD) found in human serum. Previous studies have identified that β1AR autoantibodies belongs to IgG class of immunoglobulins can modulate human β1AR responses (Iwata et al., 2001; Jahns et al., 1999a; Jahns et al., 1999b; Kaya et al., 2012). However, the use of the IgG class of β1AR autoantibodies isolated from patients with DCM showed varied responses in terms of their ability to modulate β1AR internalization and cAMP generation (Bornholz et al., 2013). The IgG class of immunoglobulins are further divided into four sub-classes namely, IgG1, IgG2, IgG3 and IgG4 (Schur, 1988; Vidarsson et al., 2014). Thus, the variability in the responses observed to IgG isolates from different patients could potentially be due to changes in relative abundance of these subclasses which may differentially alter β1AR responses and cardiac pathogenic outcomes. Consistent with this concept, recent studies show that patients who harbor the subclass of IgG3 enriched β1AR autoantibodies have significant myocardial recovery compared to the non-IgG3(+) patients (Nagatomo et al., 2017). This suggests that heart failure progression and exacerbation in the patients may depend on the representation of the IgG subclass of β1AR autoantibody.

Previous cellular studies show that pre-treatment with many of the β1AR autoantibody IgG isolates from patients positively modulates agonist isoproterenol (ISO) coupling as measured by cAMP (Bornholz et al., 2013). This suggests that the β1AR autoantibody is able to mediate a receptor conformation that allows for elevated generation of cAMP beyond the levels generated by ISO alone (Bornholz et al., 2013). This supported the concept that IgG class of β1AR autoantibodies may lead to chronic elevation in cAMP levels due to hyper-sympathetic state exacerbating the heart failure outcomes (Iwata et al., 2001; Jahns et al., 1999b; Kaya et al., 2012). In contrast, our studies show that sera containing IgG(+) β1AR autoantibodies or affinity purified IgG3(+) β1AR autoantibodies impair agonist-mediated activation β1ARs as measured by cAMP generation and adenylyl cyclase activity. This suggests that presence of the IgG3 subclass of β1AR autoantibodies may chronically reduce cAMP levels that underlie beneficial outcomes of myocardial recovery observed in these set of patients (Nagatomo et al., 2017). This supports the evolving idea that subclass members (IgG1, IgG2, IgG3 and IgG4) of the IgG family of β1AR autoantibodies may have differential effects on β1AR function as described by our studies on the IgG3 subclass of β1AR autoantibodies and substantiated by studies showing variable cAMP response to pan IgG β1AR autoantibodies (Bornholz et al., 2013).

β-blocker’s are known to block βAR responses to agonist and have been identified by their ability to inhibit generation of cAMP following agonist treatment (Kenakin, 2004; Wisler et al., 2007). Since autoantibodies mediate a β1AR confirmation that drives the receptors to generate cAMP in response to agonist (Bornholz et al., 2013), we used metoprolol, a β1AR selective blocker (antagonist) (Bristow, 1997; Prakash and Markham, 2000). It has been recognized that β-blockers alprenolol or carvedilol can bias the downstream β1AR signaling towards G-protein independent ERK activation, while simultaneously block G-protein coupling (Kim et al., 2008; Shenoy et al., 2006; Wisler et al., 2007). While metoprolol by itself blocks G-protein coupling, the presence of IgG3(+) β1AR autoantibodies surprisingly results in G-protein coupling and cAMP generation. More intriguingly, IgG3(+) β1AR autoantibodies completely abrogate agonist dobutamine mediated coupling to G-protein while remarkably restoring ERK activation. This observation suggests that IgG3(+) β1AR autoantibodies uniquely biases that agonist signaling towards G-protein independent pathway while blocking the G-protein dependent pathway.

Patients harboring the IgG class of β1AR autoantibodies are known to have worse patient outcomes (Iwata et al., 2001; Jahns et al., 1999b; Kaya et al., 2012), but the contrasting observation of benefits observed with IgG3 subclass suggests differential modulation/engagement of β1ARs to mediate beneficial signals (Nagatomo et al., 2017). The unique ability of the IgG3 subclass of β1AR autoantibodies to bias β1AR signaling through the G-protein-independent ERK pathway may underlie the beneficial outcomes in patients with this antibody (Nagatomo et al., 2017). It is known that β-blockers inhibit G-protein coupling, but also simultaneously initiates signals through β-arrestin-dependent, G-protein independent mechanisms (Kim et al., 2008; Shenoy et al., 2006; Wisler et al., 2007). Since β-blockers treatment has positive outcomes in heart failure patients, the biased G-protein-independent, β-arrestin-dependent signaling initiated by β-blockers is considered beneficial (Kim et al., 2008; Wisler et al., 2007). However, uniqueness of the IgG3(+) β1AR autoantibodies lies in their ability to modulate β1AR signaling to inhibit the classical agonist-mediated G-protein coupling, while effectively mediating G-protein-independent ERK activation in parallel. Given the hyper-sympathetic state of patients with heart failure, the presence of the IgG3(+) β1AR autoantibodies would allow for preferential engagement of the G-protein-independent pathway, reflecting in the myocardial recovery of patients harboring this sub-class of β1AR autoantibodies (Nagatomo et al., 2017). Our current studies are focused on determining whether IgG3(+) subclass of β1AR autoantibodies engages the β-arrestin-dependent EGFR transactivation pathways to mediate ERK activation while blocking G-protein coupling. Although one of the limitations of the study is the availability of IgG3(+) patient samples, we still have provided direct G-protein coupling evidence to demonstrate the distinctive ability of the IgG3(+) sub-class of the β1AR autoantibody in modulating β1AR signaling [**Fig. 5**]. IgG3 subset of autoantibodies have unique roles such as when there is higher levels of circulating β-blocker, then it allows for moderate coupling to G-proteins and when there is higher levels of circulating catecholamines, it allows for G-protein independent pathway. In summary, we present here a mechanistic pilot study that lays the foundation for: a) performing comprehensive analysis using larger cohort of patients harboring IgG3(+) β1AR autoantibodies, and b) using the subclass of IgG3(+) β1AR autoantibody as potential therapeutic strategy as patients carrying IgG3(+) β1AR autoantibodies have significantly better outcomes than IgG3(-) patients (Nagatomo et al., 2017).

**Fig 5:**
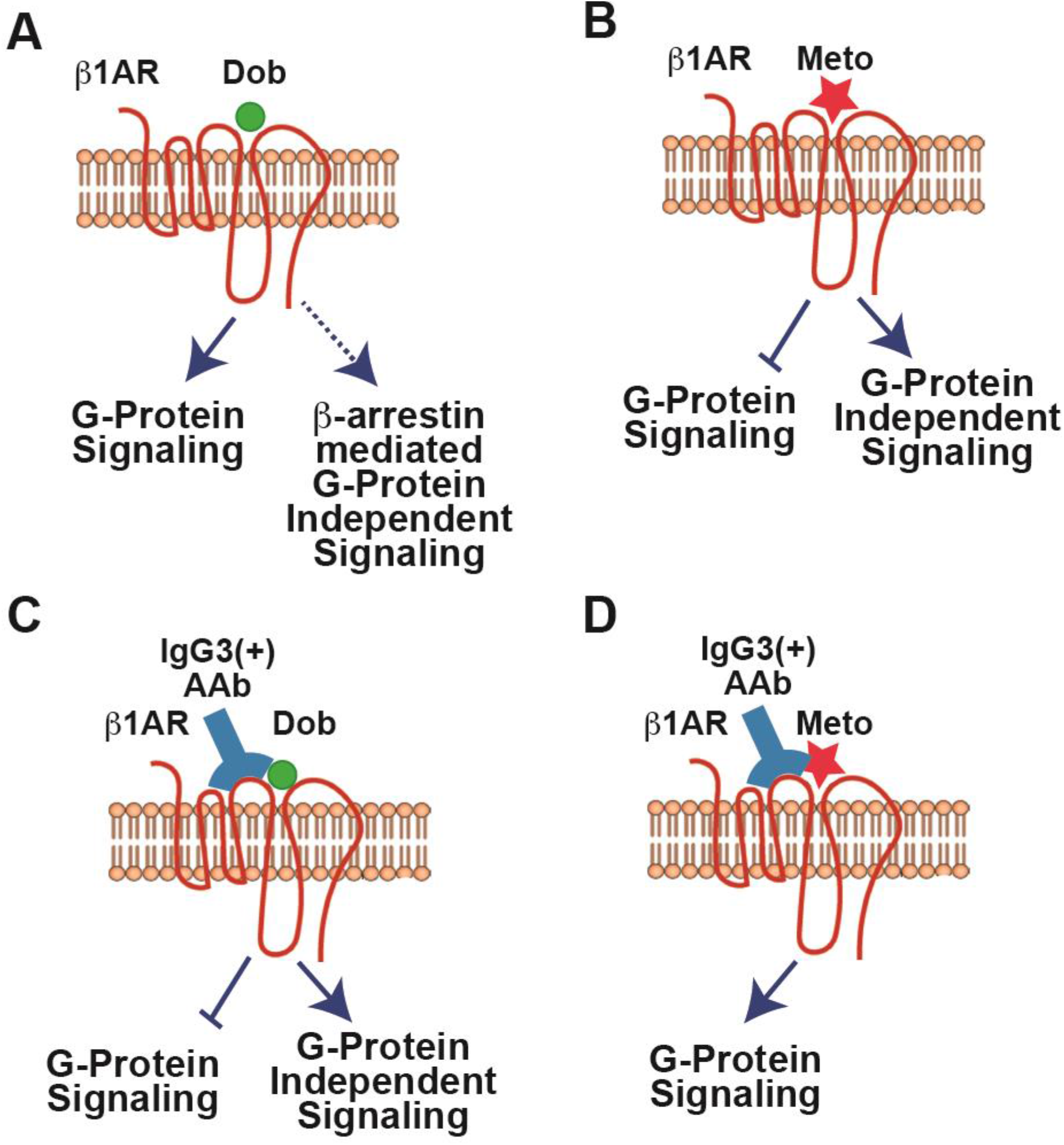
Illustration depicting signaling mechanism of IgG3(+) autoantibodies generated against human β1AR. **(A)** Agonist mediated β1AR signaling. **(B)** Antagonist mediated β1AR signaling. **(C)** Agonist mediated β1AR signaling modulated by IgG3(+) autoantibodies. **(D)** Antagonist mediated β1AR signaling modulated by IgG3(+) autoantibodies.

## MATERIALS AND METHODS

### Stable cell lines

HEK cells stably overexpressing FLAG-β1AR (FLAG-β1AR-HEK 293) was used in the experiments. Stable cell line was developed in-house by clonal selection using G418 (Geneticin) as antibiotic selection after transfecting HEK 293 cells with mammalian expression vector plasmid [pcDNA3.1(-)] containing Flag tagged - β1AR cDNA (gift from Dr. Yang K. Xiang, UC Davis, Davis, CA). HEK 293 cells stably expressing β2AR (FLAG-β2AR-HEK 293) was a gift from Dr. Robert J. Lefkowitz, Duke University, Durham, NC (Shenoy et al., 2006).

### Cell Culture

For optimal growth of cells, Minimal Essential Media (MEM) supplemented with 10% Fetal Bovine Serum and 5 % penicillin-streptomycin was used. Cells were incubated at 37 °C and 5% CO_2_.

### Cell Treatments

The cells were serum starved with serum free MEM medium for 4 hours prior to pre-treatment and stimulation. HEK-β1AR and β2AR cells were pre-treated with IgG3(+) or (-) human serum, or affinity purified IgG3(+) or (-) immunoglobulins (autoantibodies) for 30 minutes. Following pre-treatment β1AR cells were treated with specific agonist Dob or specific β1-blocker Metoprolol. HEK-β2AR cells were treated with specific agonist Iso or specific inverse agonist ICI following pre-treatment. The whole cell extracts were prepared by lysing the cells in NP-40 lysis buffer (20 mM Tris-HCl, pH 7.4; 137 mM NaCl; 1% NP40; 20% Glycerol, 1 mM PMSF; 2 μg/mL Leupeptin and 1 μg/mL Aprotinin).

### Immunoblotting

Whole cell lysates were resolved using SDS-PAGE, immunoblotted for respective proteins. The following primary antibody were used for the study; α-Flag (1:2000), α-β-actin (Sigma-Aldrich, 1:25000), α-pERK (Cell Signaling, 1:2000), α-ERK (Cell Signaling, 1:2000), α-pβ2AR (Santacruz Biotechnology, 1:1000). Appropriate HRP conjugated secondary antibody was used and chemiluminescent signals were assessed. Densitometry analysis was performed using image J software.

### Plasma Membrane isolation

The plasma membrane was isolated using a previously described protocol (Naga Prasad et al., 2001). Briefly the treated cells were scraped using osmotic lysis buffer (5 mM Tris-HCl, pH 7.4; 5 mM EDTA; 1 mM PMSF; 2 μg/mL Leupeptin and 1 μg/mL Aprotinin) and homogenized by douncing. This was followed by toggling the samples for 10 mins. The intact cells and nuclei were removed by centrifuging these samples at 2500 rpm for 5 mins. The collected supernatant was centrifuged at 17500 rpm for 30 minutes to obtain the plasma membrane as pellet. All the above steps were performed at 4oC. The pellet was resuspended in ice cold binding buffer (75 mM Tris-HCl, pH 7.5, 2 mM EDTA, and 12.5 mM MgCl2) to perform receptor binding and adenylate cyclase assay.

### Receptor binding assay (βAR density assay)

The expression of the βAR receptors on plasma membrane was done using radioligand binding assay using βAR specific radioligand _125_I-Cyanopindolol (Ferguson et al., 1996; Naga Prasad et al., 2001). This assay was performed by incubating 20 μg of plasma membrane at 37 °C for 1 hour and non-specific binding was assessed in the presence of β1AR specific β-blocker Metoprolol.

### cAMP assay

cAMP generation assay was performed using whole cell lysates. The assay was done using the standard manufacturer’s protocol. Catchpoint cAMP fluorescent assay kit from molecular devices (San Jose, CA) was used to measure the cAMP levels (Vasudevan et al., 2013).

### Adenylate cyclase assay

G-protein coupling was measured using 20 μg of plasma membrane for adenylate cyclase assay and by measuring the generated cAMP using standard procedure as previously reported (Vasudevan et al., 2011b).

### IgG purification from plasma sample

IgG was purified using a MabTrap Kit (GE #17-1128-01), based on manufacturers protocol. The kit had a binding capacity of 25 mg IgG/mL medium. Briefly the purification began with first collecting the protein from the plasma sample. This was done by adding the collected plasma to a syringe containing 1.5 mL binding buffer (20 mM sodium phosphate, pH 7.0). A 0.45 μM filter was used to flush 1 mL of the binding buffer through the syringe to get a total volume of 2.5 mL. This was followed by purification of the obtained protein, where first the column was equilibrated with binding buffer. The samples were then slowly added to the column using a syringe at a speed of ~1 drop/2 sec – 10 sec (0.2~1 mL/min). The samples were collected and reloaded back to the column. This step was repeated 4 times. This was followed by washing the samples with binding buffer until no materials appeared in the effluent. The samples were then eluted with approximately 3-5 mL of elution buffer (100 mM glycine-HCl, pH 2.7) and eluate was neutralized in a collection tube containing 375 μL of neutralizing buffer (100 mM Tris-HCl, pH 9.0). The columns were reconditioned using 5 mL binding buffer. The final step was the enrichment of the obtained IgG. The collected samples were passed through a 10 kDa Amicon Ultra tube containing 10 mL of PBS. This was centrifuged at 4000 x g for 30 min. Finally, about 200 μL of samples from the above steps were centrifuged at 14000g for 30 mins using a 3 kDa Amicon Ultra 0.5 ml device to collect the enriched IgG.

### Statistical Analysis

All data were expressed as mean±SEM (n≥3 experiments performed under identical conditions). ANOVA was used for multiple comparisons of the data. Statistical analyses were performed using GraphPad Prism© and the significance between the treatments were determined by Student *t-test*. A p-value less than < 0.05 was considered statistically significant.

## Acknowledgements

We thank Cleveland Clinic Research Programs Committee for funding of clinical study that procured the samples. This work was supported by NIH grants R01HL103931, R01DK106000, R01HL126827 to WHWT, NIH grants R01 HL089473 to SVNP and AHA Transformational Project Award 18TPA34170554. YN was supported by Research Fellowship from Myocarditis Foundation.

---

**Fig S1:**
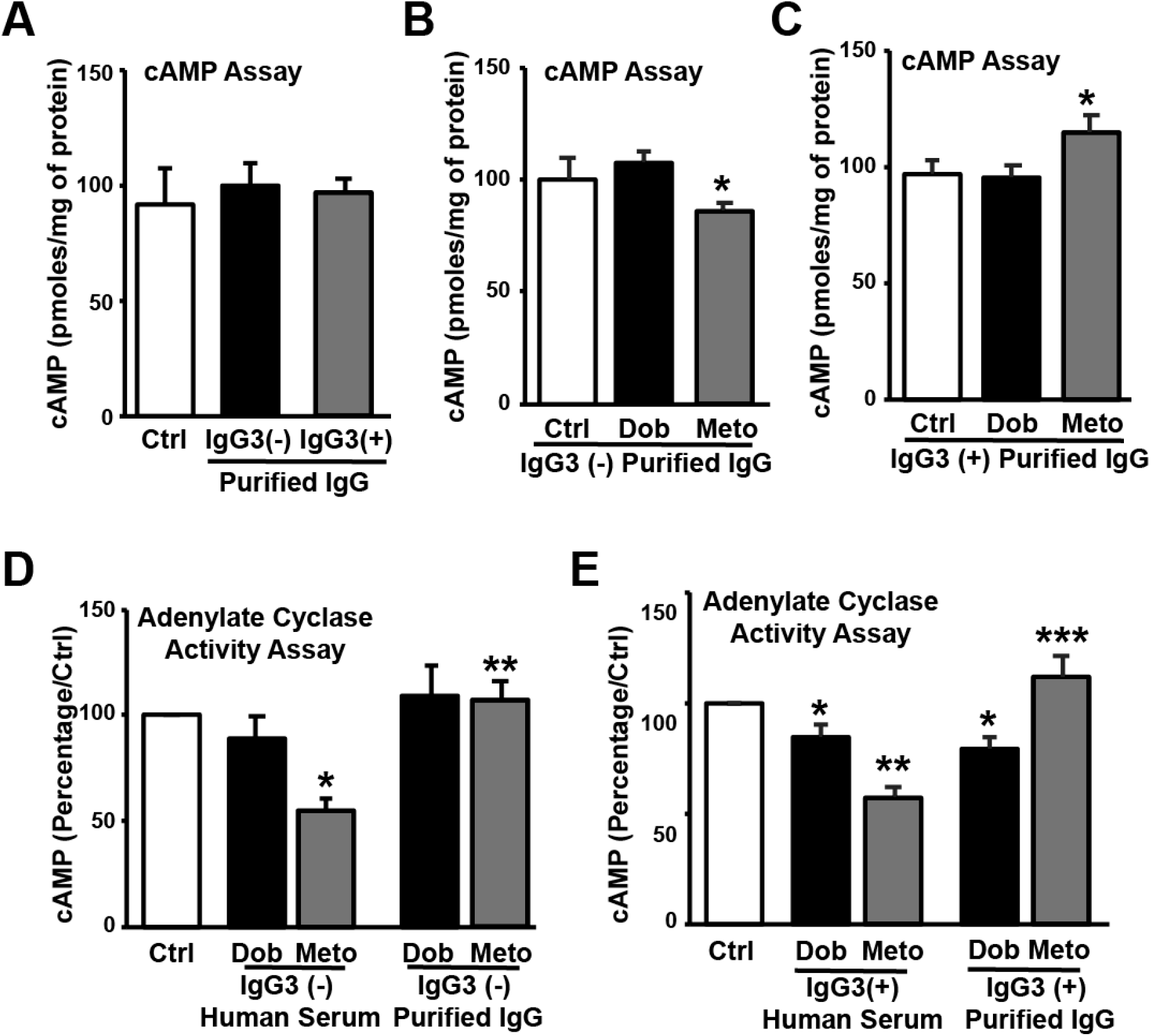
Autoantibodies (AAb) alter β1AR function. **(A)** HEK-β1AR cells were serum starved treated with affinity purified IgG3(-) and IgG3(+) AAb for 10 min. The cAMP was measured (n=3, 9). **(B)** HEK-β1AR cells were serum starved, pre-treated with affinity purified IgG3(-) AAb and stimulated with Dob or Meto. The cAMP was measured (n=9). *p ≤ 0.05 Meto vs Dob. **(C)** HEK-β1AR cells were serum starved, pre-treated with affinity purified IgG3(+) Aab and stimulated with Dob or Meto. The cAMP was measured (n=9). *p ≤ 0.01 Meto vs Ctrl/Dob. **(D)** Isolated membranes from HEK-β1AR cells were pre-treated with IgG3(-) human serum and IgG3(-) purified IgG, AC activity was performed in the presence of Dob or Meto. The amount of cAMP generated by control is expressed as 100 percent and percent change in generation of cAMP in treatments are shown (n=4, 5). *p ≤ 0.05 Meto human serum vs Ctrl/Dob human serum. **p≤ 0.001 Meto IgG3(-) purified IgG vs Meto IgG3(-) human serum. **(E)** Isolated membranes from HEK-β1AR cells were pre-treated with IgG3(+) human serum and IgG3(+) purified IgG, AC activity was performed in the presence of Dob and Meto. cAMP generated are shown as above (n=4, 7). *p ≤ 0.05 Dob human serum vs Ctrl/Meto human serum. **p ≤ 0.01 Meto human serum vs Ctrl human serum. ***p≤ 0.001 Meto IgG3(+) purified IgG vs Meto IgG3(+) human serum.

**Supplementary Fig 2:**
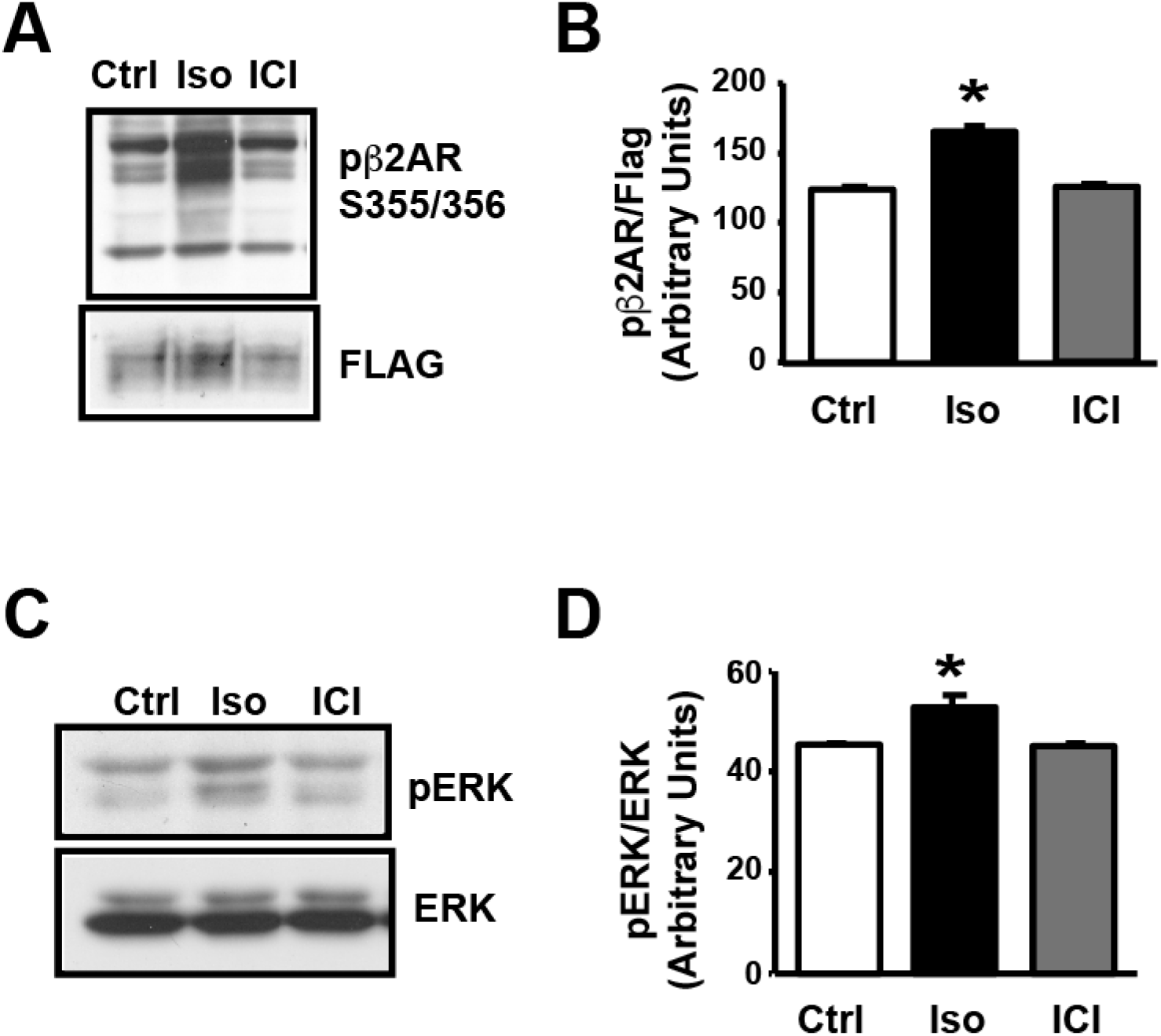
Effect of Agonist and Inverse Agonist on β2AR signaling. **(A)** HEK-β2AR cells were serum starved and stimulated with Iso or ICI. **(B)** Cumulative densitometric data for the blots (n=3). *p ≤ 0.05 Iso vs Ctrl/ICI. **(C)** HEK-β2AR cells were serum starved and stimulated with Iso and ICI, lysed and immunoblotted with anti-phospho-pERK. The blots were stripped and immunoblotted with α-ERK antibody as loading control. **(D)** Cumulative densitometric data for the blots (n=3). *p ≤ 0.05 Iso vs Ctrl/ICI.

